# MX2 viral substrate breadth and inhibitory activity are regulated by protein phosphorylation

**DOI:** 10.1101/2022.01.24.477574

**Authors:** Gilberto Betancor, Madeleine Bangham, Jun Ki Jeon, Kanisha Shah, Steven Lynham, Jose M Jimenez-Guardeño, Michael H Malim

## Abstract

Human immunodeficiency virus type-1 (HIV-1) infection is potently inhibited by human myxovirus resistance 2 (MX2/MxB), which binds to the viral capsid and blocks the nuclear import of viral DNA. We have recently shown that phosphorylation is a key regulator of MX2 antiviral activity, with phosphorylation of serine residues at positions 14, 17 and 18 repressing MX2 function. Here, we extend the study of MX2 post-translational modifications, and identify serine and threonine phosphorylation in all domains of MX2. By substituting these residues with aspartic acid or alanine, hence mimicking the presence or absence of a phosphate group, respectively, we identified key positions that control MX2 antiviral activity. Aspartic acid substitutions of residues Ser306 or Thr334 and alanine substitutions of Thr343 yielded proteins with substantially reduced antiviral activity, whereas the presence of aspartic acid at positions Ser28, Thr151 or Thr343 resulted in enhanced activity: referred to as hypermorphic mutants. In some cases, these hypermorphic mutations, particularly when paired with other MX2 mutations (e.g., S28D/T151D or T151D/T343A) acquired the capacity to inhibit HIV-1 Capsid mutants known to be insensitive to wild type MX2, such as P90A or T210K, as well as MX2-resistant retroviruses such as equine infectious anaemia virus (EIAV) and murine leukaemia virus (MLV). This work highlights the complexity and importance of MX2 phosphorylation in the regulation of antiviral activity, and in the selection of susceptible viral substrates.

**Author summary:** Productive infection by human immunodeficiency virus type-1 (HIV-1) requires the import of viral replication complexes into the nuclei of infected cells. Myxovirus resistance 2 (MX2/MxB) blocks this step, halting nuclear accumulation of viral DNA and virus replication. We recently demonstrated how phosphorylation of a stretch of three serines in the amino-terminal domain of MX2 inhibits the antiviral activity. Here, we identify additional positions in MX2 whose phosphorylation status reduces or enhances antiviral function (hypomorphic and hypermorphic variants, respectively). Importantly, hypermorphic mutant proteins not only increased inhibitory activity against wild type HIV-1, but can also exhibit antiviral capabilities against HIV-1 Capsid mutant viruses that are resistant to wild type MX2. Furthermore, some of these proteins were also able to inhibit retroviruses that are insensitive to MX2. Therefore, we propose that phosphorylation comprises a major element of MX2 regulation and substrate determination.

## Introduction

Type I interferons (IFNs), including IFN*α*, are potent inhibitors of HIV-1 infection. IFN triggers the expression of hundreds of IFN stimulated genes (ISGs), inducing an antiviral state within the host (Doyle *et al*., 2015; Bourke *et al*., 2018). One of these ISGs is myxovirus resistance 2 (*MX2/MxB*), a member of the dynamin-like family of large GTPases that also includes the closely related gene *MX1/MxA*. The MX1 and MX2 proteins have a similar domain organization, sharing 63% sequence identity (Fribourgh *et al*., 2014). They also possess antiviral activity, inhibiting a wide range of viruses, including influenza A virus (IAV), Thogoto virus (THOV) or measles virus for MX1 (Pavlovic et al., 1990; Frese et al., 1996; Kochs and Haller, 1999; Gordien et al., 2001), and HIV-1, hepatitis C virus (HCV) or herpesviruses for MX2 (Goujon *et al*., 2013; Kane *et al*., 2013; Liu *et al*., 2013; Yi *et al*., 2019; Crameri *et al*., 2018; Schilling *et al*., 2018).

The mechanisms of virus inhibition appear to be distinctive. Human MX1 requires a disordered loop (L4) for inhibition of IAV, where an interaction with the viral nucleoprotein (NP) has been shown (Mitchell et al., 2012); in contrast, the most important determinant for human MX2 inhibition is its amino-terminal domain (NTD). In particular, it has been demonstrated that transfer of this domain to otherwise non-HIV-1 inhibitory proteins (such as MX1 or Fv-1) yields proteins with antiviral activity (Goujon *et al*., 2014; Goujon *et al*., 2015). Another clear difference between MX2 and MX1 is their sub-cellular distribution, with the former localised to the cytoplasm and the nuclear envelope (Kane *et al*., 2013; Goujon *et al*., 2014; Dicks *et al*., 2018; Steiner & Pavlovic, 2020), and the latter being solely cytoplasmic (Ponten *et al*., 1997; Wisskirchen *et al*., 2011). Indeed, it has been shown that reductions in MX2 localization to the nuclear envelope may correlate with diminished antiviral activity: for instance, depletion of nucleoporin 214 (NUP214) and/or transportin 1 (TNPO1) impair the ability of MX2 to inhibit HIV-1, while also increasing cytoplasmic distribution (Dicks *et al*., 2018). Similarly, high-order oligomerization is required for MX1 inhibition of IAV (Gao *et al*., 2010), but MX2 dimers suffice to inhibit HIV-1 (Buffone *et al*., 2015; Dicks *et al*., 2015).

Early studies demonstrated that HIV-1 containing mutations in the Capsid (CA) protein can escape MX2 restriction (Goujon *et al*., 2013; Kane *et al*., 2013; Liu *et al*., 2013), pointing to CA as the determinant for MX2 sensitivity. CA proteins assemble into a lattice of hexamers and pentamers that ultimately forms the viral conical core that contains the viral genome. MX2 binds to the capsid lattice through two domains: the NTD, which is sufficient and essential for viral inhibition (Fricke *et al*., 2014; Fribourgh *et al*., 2014; Goujon *et al*., 2015; Schulte *et al*., 2015; Smaga *et al*., 2019), and the GTPase (G) domain which strengthens the association with the capsid (Betancor *et al*., 2019). However, the determinants for MX2-capsid lattice binding and viral inhibition are complex, since CA mutant viruses that are insensitive to MX2, such as N74D, P90A, G208R or T210K can still bind MX2 *in vitro* (Fricke *et al*., 2014; Fribourgh *et al*., 2014; Smaga *et al*., 2019). On the other hand, a stretch of three arginine residues at positions 11 to 13 within the MX2 NTD are essential for both viral inhibition and for capsid binding (Goujon *et al*., 2014; Fricke *et al*.; 2014; Betancor *et al*., 2019). More recently, we have shown that this triple arginine motif is required for the interaction of MX2 with myosin light chain phosphatase (MLCP), which is important for maintaining dephosphorylation at the serines at positions 14, 17 and 18 in the NTD and, as a consequence, MX2 antiviral function (Betancor *et al*., 2021).

Based on the finding that phosphorylation regulates MX2 antiviral activity, we sought to extend our assignment of MX2 phosphorylated residues, and to identify the positions of biological relevance. We obtained evidence of phosphorylation across all MX2 domains, including the GTPase catalytic site residue Thr151. We analysed the effect of phosphorylation at serine and threonine residues using phosphomimetic (aspartic acid) and non-phosphorylated (alanine) substitutions, identifying several positions where phosphorylation status altered MX2 inhibitory activity. Of note, a number of mutant proteins are described that display hypermorphic phenotypes in that they have acquired antiviral activity against MX2-insensitive CA mutant viruses, as well as against other retroviruses not inhibited by wild type MX2. These results indicate that protein phosphorylation at multiple positions in MX2 can modulate function, and suggest that post-translational modification regulates viral substrate selection.

## Results

### MX2 is phosphorylated at multiple residues, modulating antiviral activity

We have recently described a triple serine motif in the MX2 NTD (serines 14, 17 and 18) that is subjected to regulatory phosphorylation (Betancor *et al*., 2021). During the course of these analyses, we also found evidence for additional sites of phosphorylation in the NTD (*i.e*., Ser28, Thr38, Thr67 and Ser78). We therefore undertook a comprehensive analysis of MX2 to determine if residues outside the NTD are subjected to phosphorylation. To this end, MX2 was stably expressed in 293T cells, immunoprecipitated and analysed using liquid chromatography coupled with tandem mass spectrometry (LC–MS/MS). We obtained unequivocal evidence for phosphorylation at serine or threonine residues in all MX2 domains, as well as evidence for phosphorylation at other positions based on the constant neutral loss (98 Da) from the parental peptide (Fig 1, Sup Fig 1).

**Figure 1:**
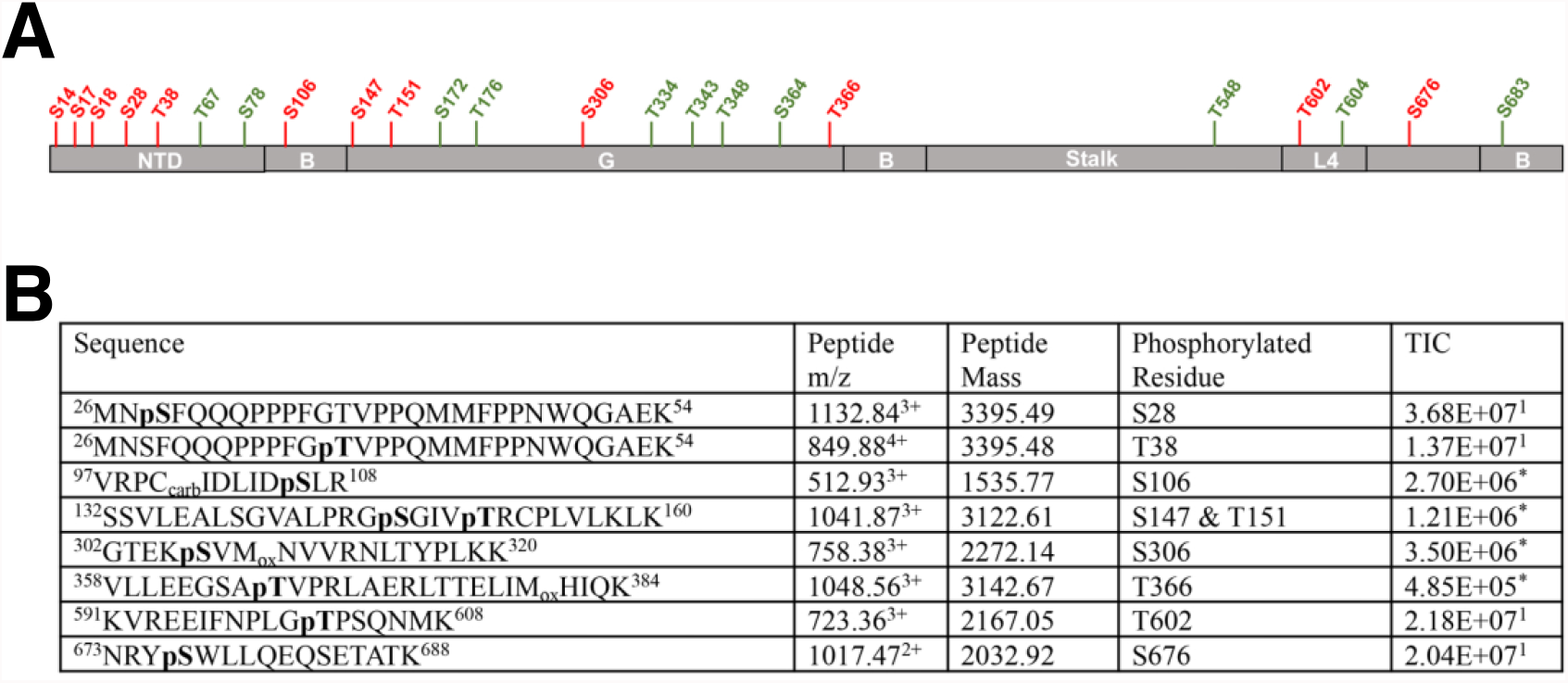
MX2 is phosphorylated at different positions. **(A)** MX2 domain organization showing the different positions found to be subjected to phosphorylation, with residues in red representing positions found to be unequivocally modified while residues in green represent positions consistent with phosphorylation. NTD, amino-terminal domain; B, bundle signalling element; G, GTPase domain; L4, Loop 4. **(B)** C-terminally Flag-tagged MX2 was expressed in 293T cells, immunoprecipitated from cell lysates, digested with trypsin and the resulting peptides subjected to MS/MS fragmentation. Obtained spectra were acquired on an Orbitrap Velos Pro and processed using Proteome Discoverer (v1.4) for identification of phosphorylated peptides. The sequences of identified peptides, specific residue(s) that are phosphorylated, their mass to charge ratio (m/z), mass, and total ion chromatogram (TIC) are shown, indicating whether the data derives from MS (^1^) or MS/MS (^*^).

We next substituted the serine or threonine residues at clear or likely sites of phosphorylation for alanine (to block phosphorylation) or aspartic acid (to mimic the presence of phosphorylation) and studied their effects on MX2 antiviral activity (Fig 2A). Most of the mutations did not impact the ability of MX2 to inhibit infection as measured by challenging U87-MG CD4/CXCR4 cells with a GFP-encoding HIV-1-based vector (HIV-1/GFP) and enumerating the number of GFP-positive cells at 48 h. We found one position which when mutated to aspartic acid produced a protein with increased antiviral activity (Ser28), two positions where mutation to aspartic acid reduced antiviral activity (Ser306 and Thr334) and two positions where HIV-1 inhibition was increased following mutation to aspartic acid, but reduced when mutated to alanine (Thr151 and Thr343). We refer to mutated MX2 proteins with enhanced antivirus function as hypermorphic mutants. Of note, three of these residues (Ser28, Thr151 and Ser306) were unequivocally identified as sites of phosphorylation in our MS/MS analysis (Fig 1, Sup Fig 1). Ser28 falls within the NTD of MX2, whereas Thr151, Ser306, Thr334 and Thr343 are all located in the GTPase (G) domain. Phosphorylation of Thr151 was particularly noteworthy, since this residue is part of the GTP binding site and its mutation to alanine has previously been described to block GTP hydrolysis (Mitchell *et al*., 2012).

**Figure 2:**
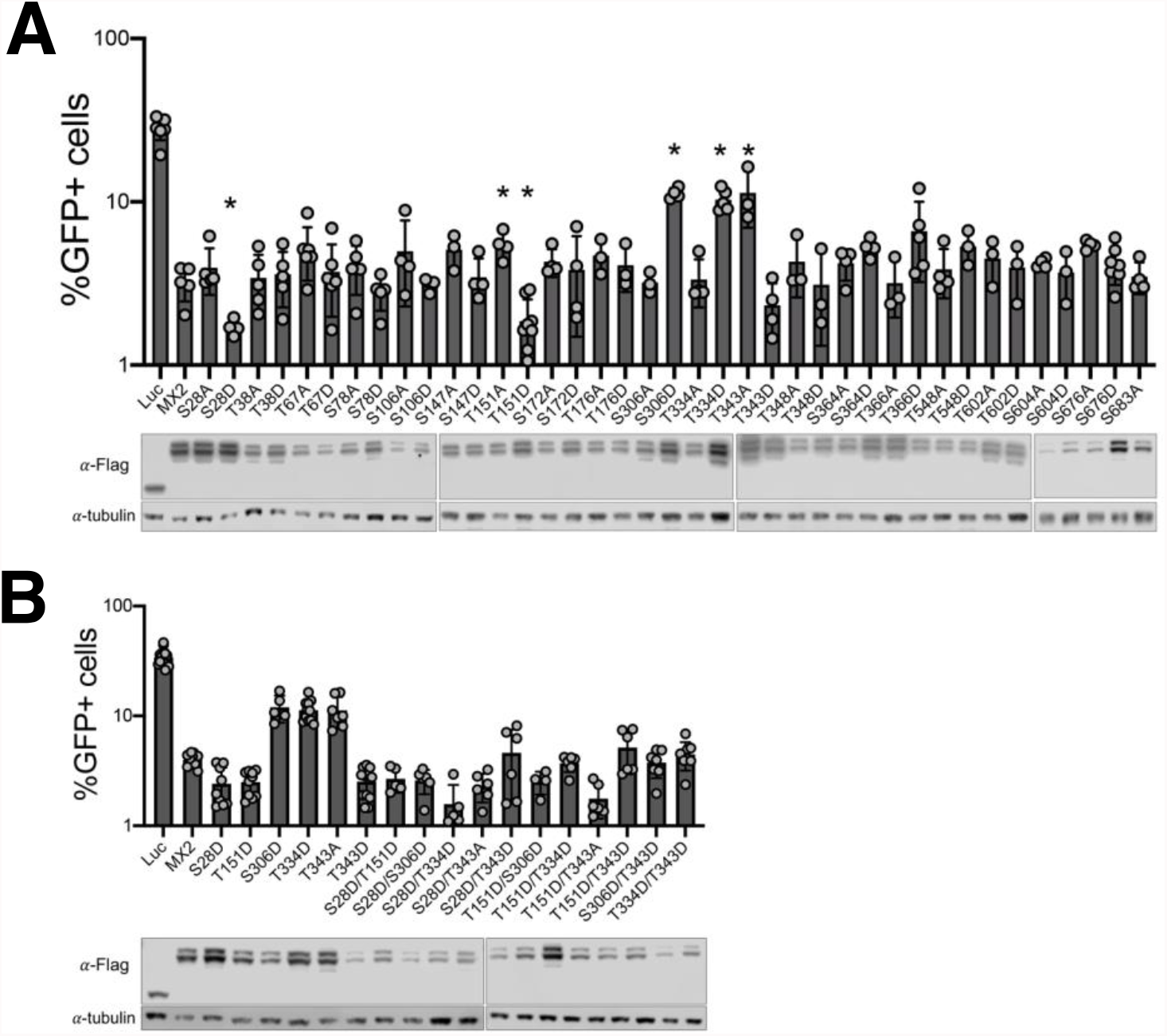
MX2 antiviral activity is modulated by phosphorylation. **(A)** Flag-tagged luciferase and wild type or mutant MX2 proteins were expressed in U87-MG CD4/CXCR4 by transduction with pEasiLV-based vectors and doxycycline induction for 48 h. The number of GFP-postive cells was determined by flow cytometry 48 h after challenging with HIV-1/GFP (n = at least 3, mean ± s.d. * = p<0.05 compared to wild type MX2, unpaired t-test). The anti-Flag immunoblot shows MX2 expression levels, with tubulin as the loading control. **(B)** U87-MG CD4/CXCR4 cells expressing MX2 proteins modified at a single or 2 positions were challenged with HIV-1/GFP, and the percentage of GFP-positive cells enumerated by flow cytometry (n = at least 5, mean ± s.d. *=p<0.05 compared to wild type MX2, unpaired t-test). The anti-Flag immunoblot shows MX2 expression levels, with tubulin as the loading control.

We explored potential patterns of dominance amongst these mutations by constructing a series of combination mutants where two positions were simultaneously mutated to aspartic acid and/or alanine. Using HIV-1/GFP as the indicator virus, we found, in general, that aspartic acid substitutions (i.e., phosphorylation) at positions Ser28 and Thr151 yielded MX2 proteins with robust antiviral activity irrespective of alanine/aspartic acid exchanges at other residues (Fig 2B). The exception to this was when Thr343 had been exchanged for aspartic acid, where the two double mutants displayed reduced inhibitory activity compared to the S28D and T151D single mutants.

Phosphorylation of serines at positions 14, 17 and 18 has been shown to inactivate MX2 (Betancor *et al*., 2021). We were therefore interested in determining whether the S28D, T151D or T343D mutations could reverse the effect of Ser14, 17 and 18 phosphorylation. Therefore, we produced MX2 proteins bearing combinations of these mutations and found that the S14, 17-18D triple mutation was dominant, with only the addition of S28D or T151D able to impart a very modest rescue of antiviral activity (Sup Fig 2).

### MX2 phosphorylation confers antiviral activity against MX2-resistant Capsid mutant viruses

Owing to the increased HIV-1 inhibitory activity conferred by phosphorylation of specific MX2 positions, we sought to study the panel of single and double mutants from Fig 2B for their antiviral activity against HIV-1 CA mutants known to be partially or fully resistant to MX2 inhibition. We analysed the CPSF6 non-binding mutant N74D (Lee *et al*., 2010; Price *et al*., 2012), the cyclophilin A (CypA) non-binding mutant P90A (Braaten *et al*., 1996; Braaten *et al*., 1997) and the MX2-resistant mutants G208R and T210K (Busnadiego *et al*., 2014) (Fig 3).

**Figure 3:**
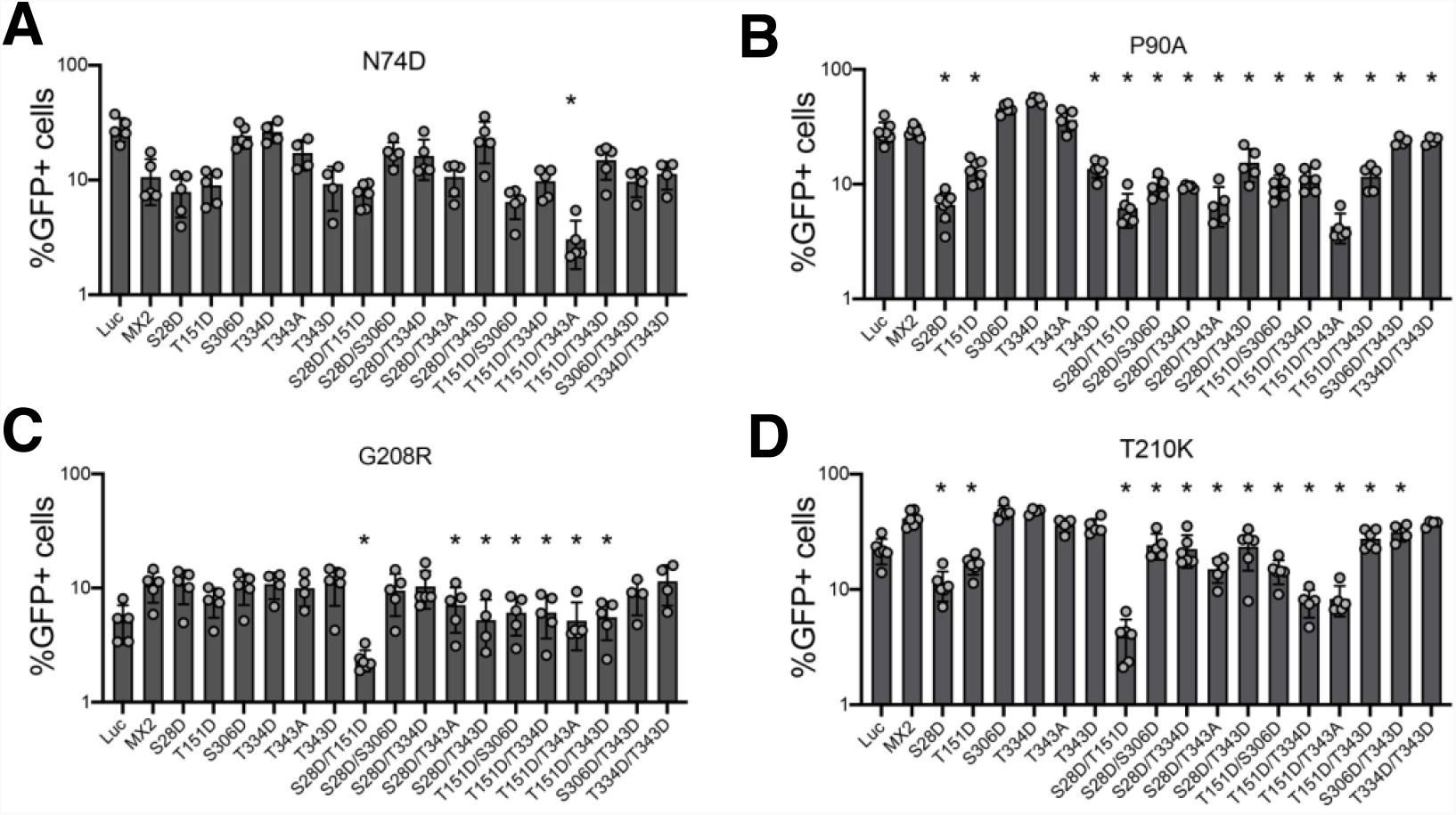
Identification of hypermorphic MX2 mutants with antiviral activity against HIV-1 Capsid mutants. U87-MG CD4/CXCR4 cells expressing wild type or mutant MX2 proteins were challenged with HIV-1/GFP bearing CA mutations N74D **(A)**, P90A **(B)**, G208R **(C)** or T210K **(D)**, and the numbers of infected cells were quantified 48 h later by flow cytometry (n = at least 5, mean ± s.d. *= p<0.05 compared to wild type MX2, unpaired t-test).

For the partially resistant N74D virus, the only mutant with a significantly enhanced antiviral phenotype was the T151D/T343A double mutant, with the remaining single and double mutants displaying either similarly weak or reduced antiviral activity compared with wild type MX2 (Fig 3A). Pro90 is located in the CypA binding loop of HIV-1 CA, and the P90A mutant is fully resistant to MX2 (Fricke *et al*., 2014; Goujon *et al*., 2014). Aspartic acid substitutions at positions Ser28, Thr151 or Thr343 alone, or when combined with other mutations, resulted in proteins with significant inhibitory activity against the P90A virus, with the S28D, S28D/T151D and T151D/T343A variants suppressing infection by 4.4-, 4.6- and 6.7-fold, respectively (Fig 3B). Gly208 and Thr210 are located at the tri-hexamer interface of the HIV-1 capsid lattice (Pornillos *et al*., 2011) and the G208R and T210K mutants are insensitive to MX2 inhibition (Busnadiego *et al*., 2014; Smaga *et al*., 2019). Several MX2 mutants demonstrated antiviral activity against these viruses, with S28D/T151D in particular inhibiting G208R and T210K viruses by 2.2- and 5.6-fold, respectively (Fig 3C and D). We interpret these results as evidence that phosphorylation of specific MX2 residues, especially Ser28 and Thr151, potentiate HIV-1 inhibitory activity not only in the context of wild type CA, but also for mutated CAs that are otherwise resistant to restriction by MX2.

### Mutant MX2 proteins can restrict non-primate retroviruses

We next asked whether MX2 mutants that inhibit MX2-resistant HIV-1 CA mutants may have extended substrate capacity and can inhibit other retroviruses not normally affected by MX2 (Goujon *et al*., 2013; Kane *et al*., 2013; Busnadiego *et al*., 2014; Matreyek *et al*., 2014). We therefore produced GFP-encoding retroviral particles based upon feline immunodeficiency virus (FIV), equine infectious anaemia virus (EIAV) or murine leukaemia virus (MLV). U87-MG CD4/CXCR4 cells expressing wild type MX2 or the S28D, T151D, T343D or S28D/T151D mutants were challenged with these stocks and infection measured at 48 h (Fig 4). Interestingly, T343D and S28D/T151D showed antiviral activity against the MX2-resistant EIAV (2.2- and 1.5-fold, respectively) (Fig 4A), and T151D, T343D and S28D/T151D were each able to inhibit MLV (1.6-, 1.5- and 1.7-fold, respectively) (Fig 4B). In contrast, none of these proteins showed any antiviral activity against FIV (Fig 4C). These results therefore reveal that phosphorylation of MX2 at key residues can broaden the profile of anti-retroviral restriction.

**Figure 4:**
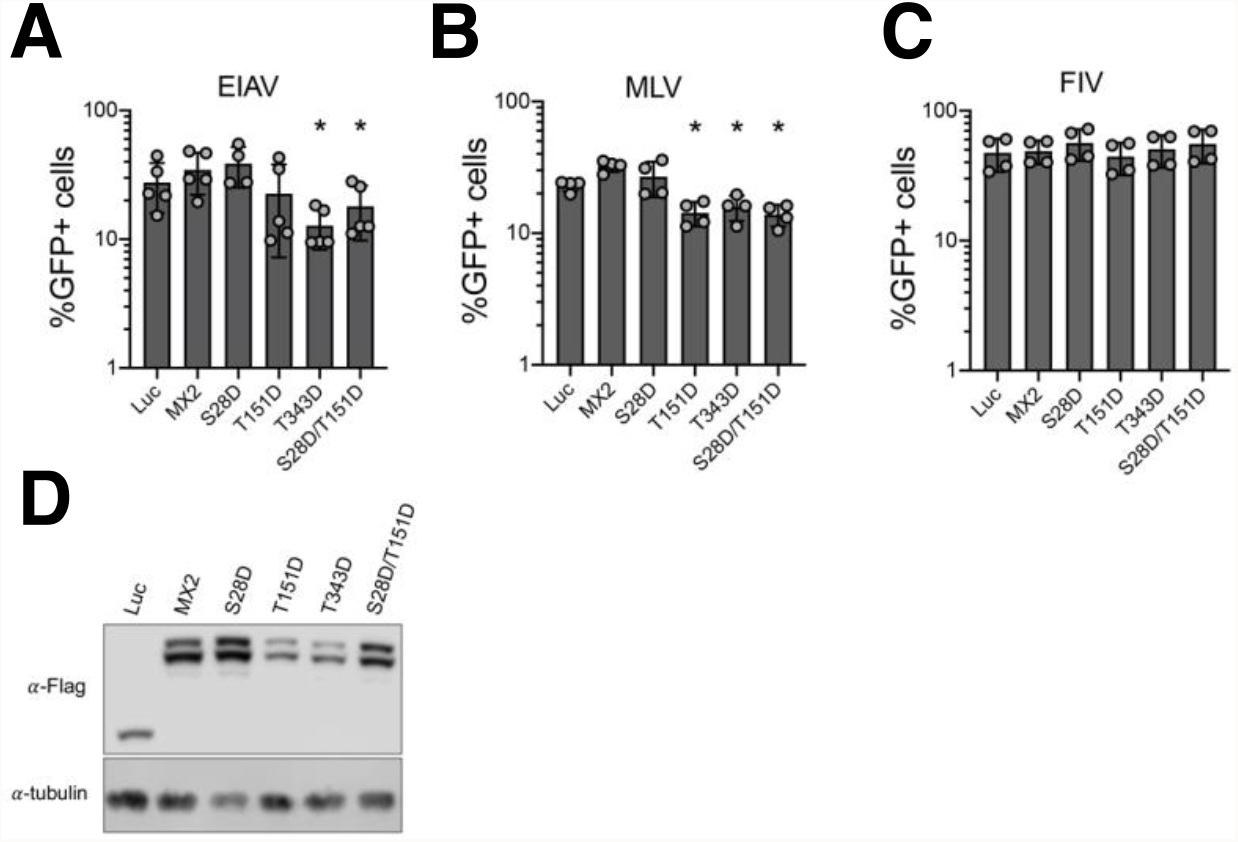
MX2 hypermorphic mutants gain antiviral activity against diverse retroviruses. U87-MG CD4/CXCR4 cells were transduced with pEasiLV expressing Flag-tagged wild type or mutant MX2 proteins. After induction for 48 h with doxycycline, cells were challenged with GFP-expressing retroviral particles based on EIAV **(A)**, MLV **(B)** or FIV **(C)**. Number of infected cells was enumerated 48 h later by flow cytometry (n = at least 4, mean ± s.d. *= p<0.05 compared to wild type MX2, unpaired t-test). **(D)** Anti-Flag immunoblot showing MX2 expression levels, with tubulin as the loading control.

### Enhanced capsid binding is not responsible for the increased antiviral activity of hypermorphic MX2 mutants

The antiviral activity of MX2 requires binding to the HIV-1 capsid through its NTD and G domains (Fricke *et al*., 2014; Fribourgh *et al*., 2014; Schulte *et al*., 2015; Smaga *et al*., 2019; Betancor *et al*., 2019; Betancor *et al*., 2021). However, capsid binding alone does not account for viral inhibition, since CA of MX2-resistant mutant viruses such as N74D, P90A or T210K readily bind MX2 (Fricke *et al*., 2014; Fribourgh *et al*., 2014; Smaga *et al*., 2019). To investigate whether the increased antiviral activity of MX2 hypermorphic mutants is linked with enhanced capsid binding, we carried out competition binding experiments using in vitro assemblies of Capsid-Nucleocapsid proteins (CANCs) (Betancor *et al*., 2019). In these experiments, Flag-tagged wild type (control) or mutant MX2 proteins were incubated with HA-tagged wild type MX2 and their ability to bind CANC complexes quantified by calculating the amount of Flag-tag protein present in the CANC-bound fraction relative to the input. We analysed the ability of MX2 mutants S28D, T151D, T343D and S28D/T151D to outcompete wild type MX2 for binding to wild type, P90A or T210K CANC complexes. While this technique is able to discern the influence of the G domain on capsid binding (Betancor *et al*., 2019), none of the hypermorphic mutants exhibited enhanced binding characteristics (Fig 5).

**Figure 5:**
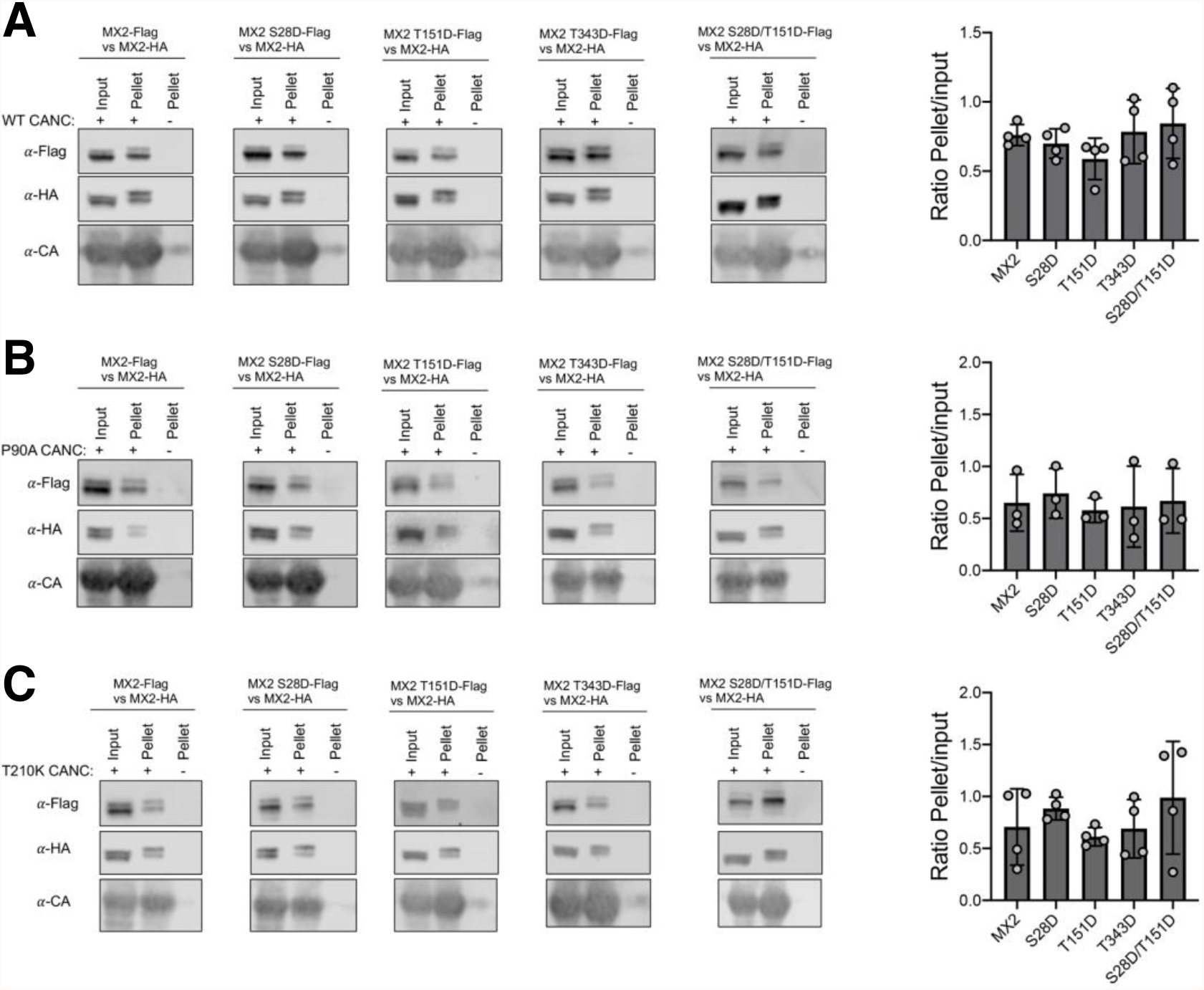
Wild type and hypermorphic MX2 mutant proteins interact with wild type or mutant HIV-1 Capsid-Nucleopcapsid nanotubes with similar affinity. Cell lysates from 293T cells expressing Flag-tagged wild type MX2, MX2 S28D, MX2 T151D, MX2 T343D or MX2 S28D/T151D were mixed with lysates from cells expressing HA-tagged wild type MX2. CANC assemblies were added to these mixtures, pelleted through a sucrose cushion, and the Flag-tagged proteins present in input and pellet fractions were quantified by immunoblot. The relative quantity of protein pelleted together with CANC complexes was inferred as the pellet/input ratio (n = at least 3, mean ± s.d).

### Nuclear envelope accumulation of MX2 hyper- and hypomorphic proteins

MX2 accumulates at the nuclear envelope mediated by a specific targeting signal in its NTD (Goujon *et al*., 2013; Kane *et al*., 2013; Schulte *et al*., 2015), and we have previously shown how diminished nuclear envelope accumulation can be associated with reduced antiviral activity (Dicks *et al*., 2018; Betancor *et al*., 2021). We therefore asked if the modified inhibitory activity of MX2 hypo- or hypermorphic proteins could be correlated with changes in their cellular distribution. We expressed wild type and mutant proteins in HeLa cells and analysed the localization of MX2 and the nuclear pore complex component NUP358 by immunofluorescent staining. The localization of MX2 to the nuclear envelope was measured as the percentage of protein colocalizing (Mander’s coefficient) with NUP358 (Fig 6, Sup Fig 3).

**Figure 6:**
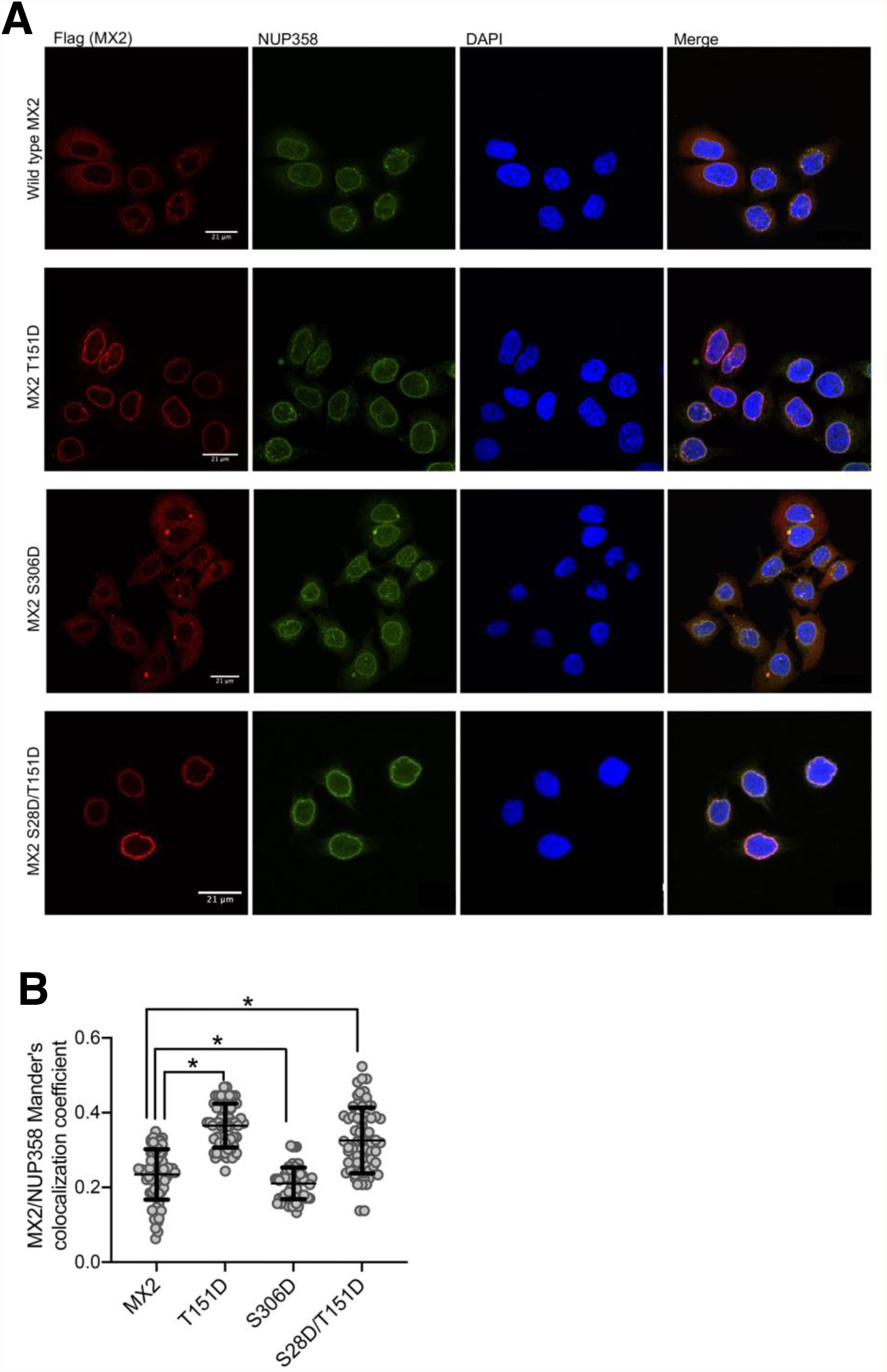
MX2 hypo- and hyper-morphic proteins with altered nuclear envelope accumulation. **(A)** Flag-tagged wild type MX2, or the T151D, S306D or S28D/T151D mutants were stably expressed in HeLa cells by transduction with puromycinR lentiviral vectors. Cells were seeded onto glass coverslips and localization of MX2 and endogenous NUP358 were determined by immunostaining with anti-Flag and anti-NUP358 antibodies and confocal microscopy. DAPI was used to stain the nuclei. **(B)** Nuclear envelope accumulation of MX2 was quantified as the colocalization of MX2 and NUP358 (Mander’s coefficient) for 70 ramdonly selected cells per condition (*=p<0.05, unpaired t-test).

To summarise, the results did not reveal a consistent correlation between increased or reduced antiviral activity and the extent of MX2 nuclear envelope accumulation. For instance, the hypomorphic mutant T334D or hypermorph variants such as S28D and T343D showed the same cellular distribution as wild type MX2 (Sup Fig 3). Nevertheless, we noted that the less active mutant S306D displayed diminished nuclear envelope localization, whereas proteins with aspartic acid at position 151 (*i.e*. T151D and S28D/T151D) showed increased in nuclear envelope accumulation (Fig 6).

## Discussion

The precise mechanism of MX2 inhibition of HIV-1 infection is currently unknown. A growing body of evidence has highlighted the central importance of the MX2’s NTD (Goujon *et al*., 2013; Kane *et al*., 2013; Liu *et al*., 2013; Goujon *et al*., 2015), oligomerization (Buffone *et al*., 2015; Dicks *et al*., 2015), binding to the viral capsid (Fricke *et al*., 2014; Fribourgh *et al*., 2014; Betancor *et al*., 2019; Smaga *et al*., 2019), interactions with cellular binding partners (Kane *et al*., 2018; Dicks *et al*., 2018, Betancor *et al*., 2021), and regulatory phosphorylation/ dephosphorylation at the serines at positions 14, 17 and 18 (Betancor *et al*., 2021).

Here, we build on this final concept, showing that MX2 is phosphorylated at multiple positions throughout the protein (Fig 1, Sup Fig 1). Using aspartic acid and alanine substitution mutants, we found that the phosphorylation status at many of these positions does not affect the antiviral activity of MX2 (Fig 2). Importantly, we did identify a number of residues whose phosphorylation status has a significant impact on MX2 function: phosphorylation of Ser28 yielded a protein with increased activity, while phosphorylation of Ser306 or Thr334 resulted in less active variants. Interestingly, we also found two positions showing increased or decreased activity depending on the nature of the introduced mutation: phosphorylation of Thr151 or Thr343 resulted in proteins with enhanced antiviral activity, whereas the corresponding dephosphorylated forms were weaker inhibitors of HIV-1 (Fig 2). Taking these data together with earlier observations that phosphorylation of serines 14, 17 and 18 decrease MX2 activity, a complex picture emerges where multiple different phosphorylation events can repress or activate MX2. We predict that multiple kinases likely phosphorylate different serines and threonines in MX2, and that their identification (and that of the counteracting phosphatases) as well as an appreciation of their regulatory mechanisms will be required to understand the post-translational regulation of MX2 antiviral activity.

Phosphorylation of Thr151 is especially interesting because this residue is part of the MX2 GTPase catalytic site and is directly involved in GTP hydrolysis (Mitchell *et al*., 2012; Fribourgh *et al*., 2014). Mutant T151A has been presumed to be unable to hydrolyse GTP (King *et al*., 2004), while maintaining antiviral activity. This observation has been used as evidence that GTPase catalytic activity is dispensable for HIV-1 inhibition (Matreyek *et al*., 2014). However, recent findings have highlighted the importance of the GTPase domain for specific attributes of MX2, including protein oligomerization (Alvarez *et al*., 2017) and HIV-1 capsid binding (Betancor *et al*., 2019). We anticipate that T151D is unable to bind GTP, due to the steric hindrance between GTP and the negative charge provided by the introduced aspartic acid. However, in contrast to T151A, the antiviral activity of T151D is enhanced, and we hypothesize that this could be due to the negative charge of Asp151 in the GTP binding site mimicking that of occupancy by the phosphate groups of GTP.

The ability of the majority of the phosphorylated or non-phosphorylated variant proteins to accumulate at the nuclear envelope was indistinguishable from the wild type protein. Exceptions to this were the less active S306D mutant which presented with a more diffuse distribution, and the T151D and S28D/T151D proteins which localized more intensely at the nuclear envelope (Fig 6). We have previously linked diminished nuclear envelope accumulation with reduced antiviral activity; for instance, depletion of NUP214 and/or TNPO1, or phosphorylation of the NTD triple serine motif S14, 17-18, favoured a more diffuse cytoplasmic distribution together with reduced anti-HIV-1 activity (Dicks *et al*., 2018; Betancor *et al*., 2021). However, through the analysis of many mutated proteins herein, this relationship is evidently not always observed: the S28D, T151A, T334D and T343A variants demonstrate this clearly with significant alterations in their antiviral activity failing to correlate with measurable changes in nuclear envelope accumulation. As such, these changes in antiviral phenotypes will need to be ascribed to mechanism(s) distinct from altered targeting to the nuclear envelope.

Since we identified MX2 mutants with increased anti-HIV-1 activity, we also tested their ability to inhibit known MX2-resistant HIV-1 mutants (Fig 3). While the N74D CA mutant largely retained resistance to all MX2 forms, with only the T151D/T343A compound mutation significantly improving activity, we found several variants that can inhibit P90A, G208R or T210K CA mutant viruses. Most interesting are S28D, T151D and S28D/T151D, which strongly inhibited all three viruses. However, upon examining the in vitro capsid (wild type, P90A and T210K) binding ability of these mutants, we were unable to discern any significant changes (Fig 5). When considered alongside the established binding of wild type MX2 to non-restricted capsids such as P90A and T210K (Fricke *et al*., 2014; Smaga *et al*., 2019), we conclude that CANC interaction assays do not accurately predict patterns of MX2-mediated restriction. Indeed, capsid binding is known to be complex, with several MX2/CA surfaces involved (Fricke *et al*., 2014; Fribourgh *et al*., 2014; Betancor *et al*., 2019; Smaga *et al*., 2019), and alternative and perhaps more discriminatory approaches will need to be employed/developed to determine whether differences in the extent or mode of binding of mutant MX2 or CA proteins can help explain profiles of restriction.

Finally, we tested the capacity of MX2 hypermorphic proteins to inhibit other retroviruses. In agreement with previous data, MX2 did not inhibit FIV, EIAV or MLV replication (Fig 4) (Goujon *et al*., 2013; Busnadiego *et al*., 2014; Matreyek *et al*., 2014), and while FIV was resistant to all proteins tested, phosphorylation of positions Thr343 (for MLV), Ser28 (for EIAV and MLV) and Ser28/Thr151 combined (for EIAV and MLV) yielded MX2 variants with restriction activity. While the extents of inhibition are relatively weak (i.e., 1.5- and 2.2-fold), these findings raise the interesting possibility of post-translational modifications being able to modulate the spectrum of MX2 substrate viruses. With the range of human viruses identified as sensitive to human MX2 growing (now including hepatitis B and C viruses and herpesviruses (Wang *et al*., 2020; Yi *et al*., 2019; Staeheli *et al*., 2018; Schilling *et al*., 2018; Crameri *et al*., 2018)), we suggest that screens with MX2 mutants that represent the phosphorylated or non-phosphorylated state at different residues may facilitate the identification of additional MX2-susceptible viruses.

In summary, while important questions remain to be answered regarding the antiviral activities of MX proteins, not the least the identification of the regulatory kinase(s) and phosphatase(s), this study highlights the importance of post-translational modification in modulating MX2 function.

## Material & Methods

### Retroviral vectors

GFP-expressing retroviral particles based on murine leukemia virus (MLV/GFP), feline immunodeficiency virus (FIV/GFP), equine infectious anaemia virus (EIAV/GFP), wild type, N74D-, P90A-, G208R- or T210K-Capsid (CA)-bearing HIV-1 (HIV-1/GFP), as well as the IRES-puromycin N-acetyltransferase (puromycinR) lentiviral vector and the pEasiLV inducible vector systems have been described (Goujon *et al*., 2013; Bulli *et al*., 2016; Dicks *et al*., 2015; Dicks *et al*., 2018). In brief, particles were produced by co-transfection of 293T cells with the specific Gag-Pol plasmid, a vesicular stomatitis virus G (VSV-G) envelope expression vector (pMD.G) and the specific retroviral vector at a ratio of 1:0.25:1, respectively. In all cases, transfections were performed using TransIT-2020 (Mirus) reagent at a 2:1 ratio with DNA. 24 h after transfection, culture medium was changed. 48 h after transfection the medium was filtered through 0.45 μm cut-off filters (Sartorius) to purify viral particles. Resulting stocks were either stored at −80 °C for future use (MLV/GFP, HIV-1/GFP, HIV-2/GFP, SIV/GFP, FIV/GFP and EIAV/GFP) or used immediately for transductions (puromycinR and pEasiLV).

### Cells

293T, HeLa and parental U87-MG cells were obtained from the American Tissue Culture Collection (ATCC). The generation of the U87-MG CD4+ CXCR4+ cell line has been described (Goujon *et al*., 2013). All cells were cultured in Dulbecco’s modified Eagle medium (DMEM) supplemented with heat-inactivated fetal bovine serum (10%), L-glutamine, penicillin (100 U/ml) and streptomycin (100 mg/ml).

### Plasmid Constructs

MX2 cDNA tagged with a Flag tag was cloned into pCAGGs (Adgene), pEasiLV and/or the puromycinR plasmid. MX2 mutants were produced by site directed mutagenesis using standard PCR-based methods. *Photinus pyralis* luciferase-Flag was cloned into pEasiLV. Wild type HIV-1 CANC (Gag residues 133–432), was cloned into pET-11a and the N74D, P90A, G208R and T210K mutants derived by site directed mutagenesis.

### Retroviral vector infectivity assays

U87-MG CD4/CXCR4 cells were transduced with pEasiLV-based vectors as described (Goujon *et al*., 2014). Briefly, 8 h after transduction, culture medium was replaced with fresh medium containing 0.5 µg/ml doxycycline to induced protein expression. 40 h later, cells were challenged with retroviral vectors, and 48 h later the percentage of cells expressing GFP was enumerated by flow cytometry (FACSCanto II; BD Biosciences).

### CANC pull-down competition experiments

Wild type and mutant CANC proteins were expressed in Escherichia coli Rosetta (DE3) cells (Merck Millipore) as previously described (Betancor *et al*., 2019). For their assembly, CANC complexes were diluted to 40 μM final concentration in 50 mM Tris-HCl pH 8 containing 100 mM NaCl and 5 μM of TG50 oligonucleotide (5′-TG(25)-3′) and incubated overnight at room temperature. HA-tagged wild type MX2 and Flag-tagged protein overexpression was achieved by transient transfection of 293T cells. 48 h after transfection, cells were recovered and lysed in 500 µl hypotonic lysis buffer (10 mM Tris-HCl pH 8, 10 mM KCl, 1× complete protease inhibitor cocktail (Roche)), using a Dounce homogeneizer. The resulting lysate was centrifuged at 20,000 x g for 15 min at 4 °C to clear cell debris. Then, 200 µl of MX2-HA lysate was mixed with 200 µl of the corresponding Flag-tag MX2 proteins and with either 40 μl of 40 μM assembled CANC or with 40 μl of capsid assembly buffer (50 mM Tris-HCl pH 8, 100 mM NaCl) containing 5 μM TG50 (as a control); an input sample was withdrawn from the former. Samples were incubated for 1 h at room temperature, overlaid on 250 μl of 70% sucrose cushion and centrifuged at 20,000 x g for 15 min at room temperature. The supernatant was removed, and the pellet washed with 500 μl of wash buffer (50 mM Tris-HCl pH 8, 50 mM NaCl and 5 mM KCl) before being re-pelleted at 10,000 x g for 8 min. Finally, the resulting pellet was resuspended in 50 μl sample buffer. The Input and Pellet fractions (with or without CANC) were analysed by immunoblotting using anti-Flag monoclonal M2 (Sigma, F3165), HRP-conjugated anti-HA (Sigma, 3F10) and anti-CA (Fouchier *et al*., 1997) antibodies.

### Immunoblots

pEasiLV transduced U87-MG CD4/CXCR4 cells were lysed in lysis buffer (10 mM Tris-HCl pH 8,150 mM NaCl, 1 mM EDTA, 1% Triton X-100 and 0.1% sodium deoxycholate). Lysates were briefly sonicated, cleared by centrifugation at 1,500 x g for 10 min at 4 °C, and boiled for 10 min before resolving in 12% SDS–polyacrylamide gels. Proteins were electrophoretically transferred to a 0.45 μm nitrocellulose membrane (Amersham) and their presence was analysed using HRP-conjugated anti-Flag (Sigma, A8592) and anti-α-tubulin (Abcam, ab15246) antibodies.

### Fluorescence microscopy

HeLa cells were transduced with puromycinR constructs bearing Flag-tagged wild type or mutant MX2 cDNAs. 48 h after transduction, stably expressing cells were selected with 1 μg/ml puromycin for 72 h and seeded onto coverslips at ∼50,000 cells per well in 24-well plates. 24 h later, cells were washed with 1x phosphate buffer saline and fixed in 4% paraformaldehyde (EM Sciences) for 15 min. Cell membrane permeabilization was achieved by incubation with 0.2% Triton X-100 for 15 min. Then, the coverslips were blocked with NGB buffer (50 mM NH4Cl, 2% goat serum, 2% bovine serum albumin) for 1 h. MX2-Flag proteins were detected using anti-Flag monoclonal M2 and secondary antibody conjugated to Alexa 594 (Invitrogen, A21203). Endogenous NUP358 was detected using rabbit anti-NUP358 antibody (Abcam, ab64276) and secondary antibody conjugated to Alexa 488 (Invitrogen, A21206). The nuclei were labelled using DAPI (4’,6-diamidino-2-phenylindole) staining (0.1 mg/ml for 5 min). Cells were visualized using a Nikon A1 point-scanning laser confocal microscope (Nikon Instruments). Quantification of the nuclear envelope accumulation of wild type or mutant MX2 proteins was calculated as the percentage of total protein colocalizing with NUP358, using Manders’ coefficient.

### MX2 phosphorylation and mass spectrometry

293T cells stably expressing MX2-Flag were lysed in PBS containing 1% Triton-X100, supplemented with 1× PhosSTOP phosphatase inhibitor cocktail (Sigma) and 1× complete protease inhibitor cocktail (Roche). The lysate was then centrifuged at 16,000 x g for 10 min, and the resulting supernatant incubated with anti-Flag magnetic beads (Sigma) for 2 h at 4 °C. The beads were washed 3 times with lysis buffer, and MX2-Flag was eluted by resuspending the beads in sample buffer and boiled for 10 min. Immunoprecipitated MX2 was resolved on a 12% SDS–polyacrylamide gel (Thermo Fisher), stained with SimplyBlue (Thermo Fisher) and the band corresponding to MX2 was excised and subjected to in-gel reduction (dithiothreitol; Sigma), alkylation (Iodoacetamide; Sigma) and trypsin (bovine; Sigma) digestion. Peptides were extracted from the gel by a series of washes of acetonitrile and water and subsequently lyophilised. Resulting samples were resuspended in 50 mM NH_4_HCO_3_ and analysed by LC–MS/MS. Chromatographic separations were performed using an EASY nLC-II nanoflow system (Thermo Fisher Scientific). Reversed-phase chromatography was used to resolve peptides using a 10 cm 75 μm C18 PepMap column and three-step linear gradient of MeOH (100%) in CH_2_O_2_ (0.1%) at 300 nl min−1 with peptides resolved using a gradient rising from 5 to 40% solvent B (0.1% formic acid in 100% acetonitrile) by 180 min. The peptides were ionized by electrospray ionization at 2.6 kV using a stainless-steel emitter at 250°C. All spectra were acquired using an Orbitrap Velos Pro (Thermo Fisher Scientific) operating under Xcalibur v2.2 operated in data-dependent acquisition mode. FTMS1 spectra were collected at a resolution of 60,000 FWHM (full width half maximum of the peak) over a scan range (m/z) of 400-1800. Automatic gain control (AGC) was set at 1×106 with a maximum injection time of 50 ms (3 sec duty cycle). ITMS2 spectra were collected with a maximum AGC target of 3×104, maximum injection time 50 ms and CID collision energy of 35%.

Data obtained were processed in Proteome Discoverer (PD, Thermo Fisher Scientific) v1.4 and interrogated using Mascot (v1.5) with an in-house curated database including the sequence of the MX2 protein (P20592). Variable modifications included carbamidomethylation (Cys), oxidation (Met) and phosphorylation (Ser, Thr & Tyr) at 5% FDR for protein and peptide tolerance of 20 ppm. MSF files generated by PD were uploaded into Scaffold software (Proteome Software Inc, v5.0) for alignment, visualisation and manual phospho-PTM verification.

## Acknowledgements

We thank Stelios Papaioannou and Adrian Signell for the provision of reagents and helpful discussions. For the purpose of open access, the author has applied a CC BY public copyright license to any Author Accepted Manuscript version arising from this submission.

