## Supplementary figures 1 to 3 for "MX2 viral substrate breadth and inhibitory activity are regulated by protein phosphorylation"

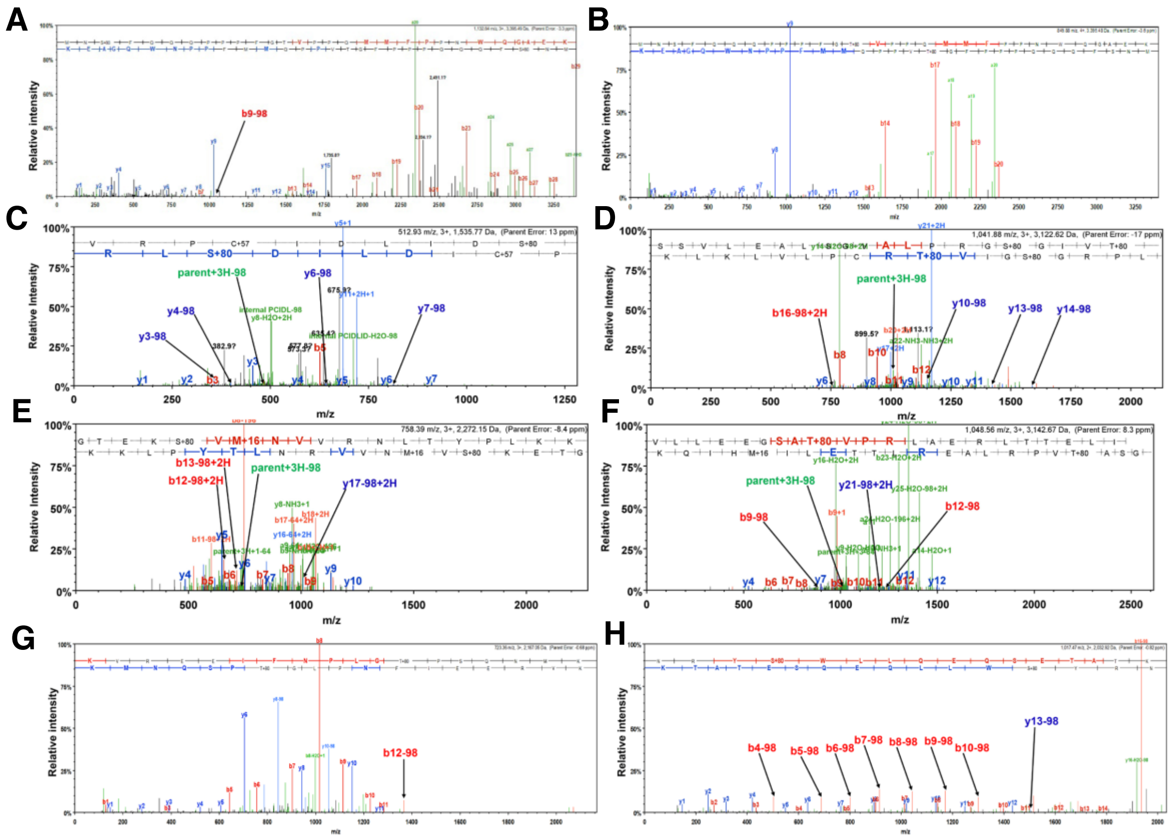


**Supplementary figure 1: Phosphorylation of MX2.** LC-MS/MS data generated from trypsin digested MX2 peptides modified by phosphorylation. MS/MS fragmentation spectra were acquired on an Orbitrap Velos Pro and processed using Proteome Discoverer (v1.4) and visualised in Scaffold software. Fragmentation patterns following database assignment show y-ions from the peptide C-terminal side and b-ions from the N-terminal side along with loss of $H_2O$ and NH3. Peaks assigned with -98 Da represent the neutral loss of phosphoric acid from the modified residue and the subsequent residues in the peptide sequence determining correct site localisation. **(A)** Ser28, **(B)** Thr38, **(C)** Ser106, **(D)** Ser147 and Thr151, **(E)** Ser306, **(F)** Thr366, **(G)** Thr602, **(H)** Ser676.

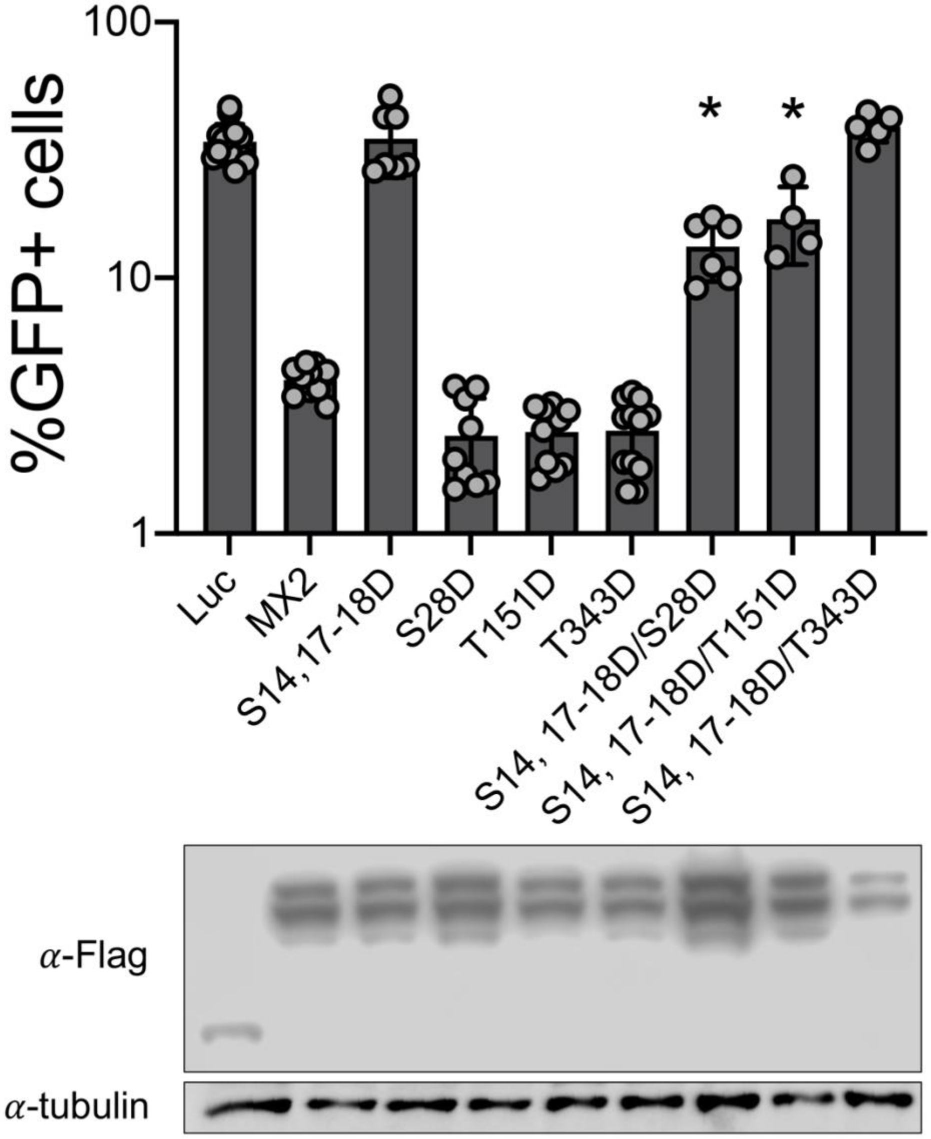


**Supplementary figure 2: Phosphorylation of MX2 triple serine motif at positions 14, 17 and 18 severely reduced the antiviral activity of hypermorph proteins.** U87-MG CD4/CXCR4 cells expressing Flag-tagged wild type and mutant MX2 proteins were challenged with HIV-1/GFP particles, and the number of infected cells enumerated 48 h later by flow cytometry (n = at least 4, mean ± s.d. *=p<0.05 compared to luciferase (Luc), unpaired t-test). Bottom, anti-Flag immunoblot shows MX2 expression levels, with tubulin as the loading control.

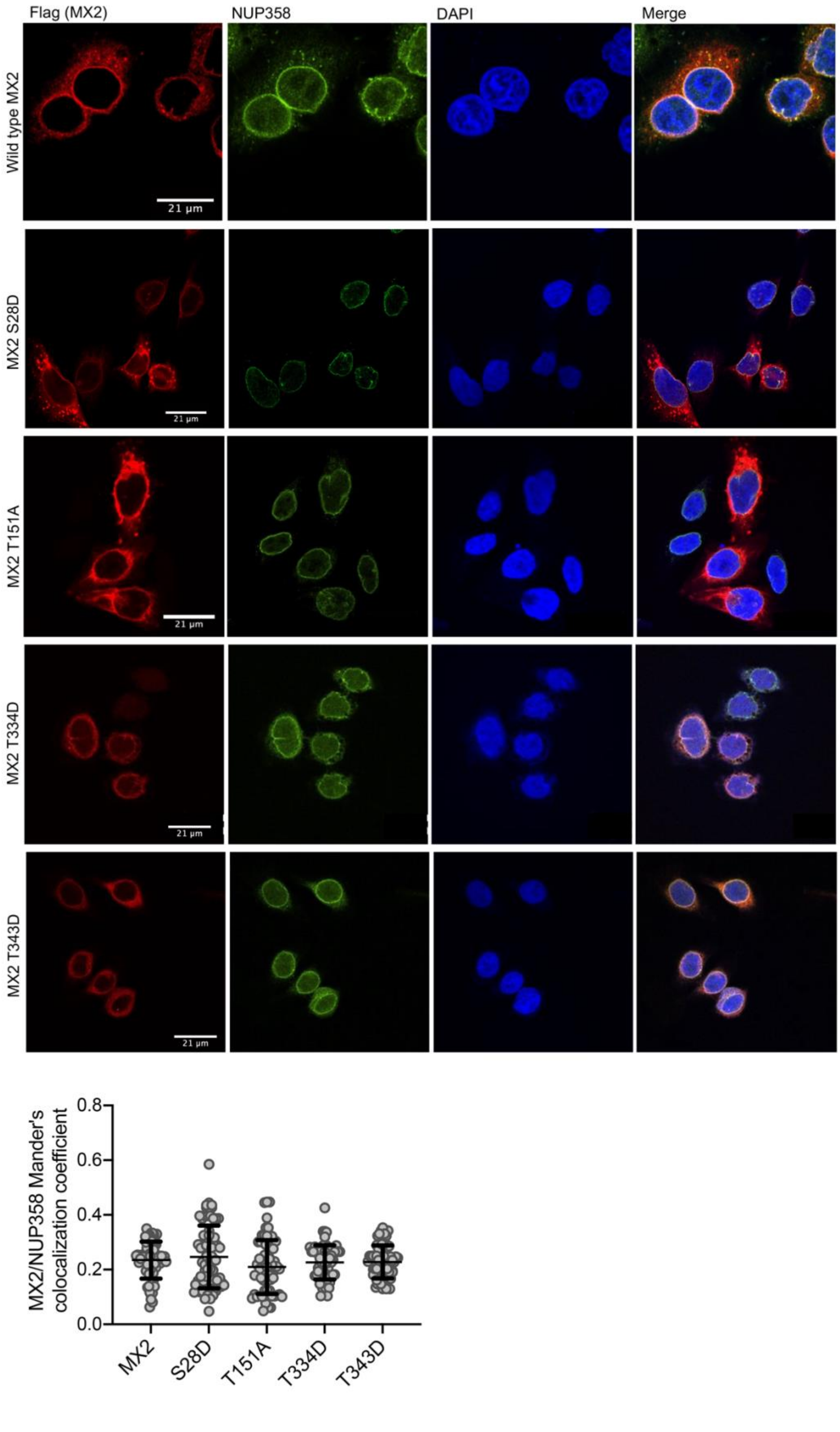

Flag (MX2)
NUP358
DAPI
Merge
Wild type MX2
MX2 S28D
MX2 T151A
MX2 T334D
MX2 T343D
21 µm
MX2/NUP358 Mander's colocalization coefficient
0.8
0.6
0.4
0.2
0.0
MX2
S28D
T151A
T334D
T343D

**Supplementary figure 3: MX2 hypo- and hyper-morphic proteins with unaltered nuclear envelope accumulation.** Hela cells stably expressing Flag-tagged MX2 proteins were seeded in coverslips and immunostained to detect the presence of the Flag peptide and endogenous NUP358, with Dapi serving to mark the nuclei. MX2 colocalization with NUP358 (Mander's coefficient) was used to quantify its nuclear envelope accumulation (n=70, *=p<0.05, unpaired t-test).
